# Reporting of “dialysis adequacy” as an outcome in randomised trials conducted in adults on haemodialysis: a systematic review

**DOI:** 10.1101/453191

**Authors:** Sanne Steyaert, Els Holvoet, Evi Nagler, Simon Malfait, Wim Van Biesen

## Abstract

**Background:** Clinical trials are most informative for evidence-based decision-making when they consistently measure and report outcomes of relevance to stakeholders, especially patients, clinicians, and policy makers. However, sometimes terminology used is interpreted differently by different stakeholders, which might lead to confusion during shared decision making. The construct *dialysis adequacy* is frequently used, suggesting it is an important outcome both for health care professionals as for patients.

**Objective:** To assess the scope and consistency of the construct *dialysis adequacy* as reported in randomised controlled trials in hemodialysis, and evaluate whether these align to the insights and understanding of this construct by patients.

**Methods:** To assess scope and consistency of *dialysis adequacy* by professionals, we performed a systematic review searching the Cochrane Central Register of Controlled Trials (CENTRAL) up to July 2017. We identified all randomised controlled trails (RCT) including patients on hemodialysis and reporting *dialysis adequacy*, *adequacy* or *adequacy of dialysis* and extracted and classified all reported outcomes. To explore interpretation and meaning of the construct of *adequacy* by patients, we conducted 10 semi-structured interviews with HD patients using thematic analysis. Belgian registration number B670201731001.

**Findings:** From the 31 included trials, we extracted and classified 98 outcome measures defined by the authors as *adequacy of dialysis*, of which 94 (95%) were biochemical, 3 (3%) non-biochemical surrogate and 2 (2%) patient-relevant. The three most commonly reported measures were all biochemical. None of the studies defined *adequacy of dialysis* as a patient relevant outcome such as survival or quality of life.

Patients had a substantially different understanding of the construct *dialysis adequacy* than the biochemical interpretation reported in the literature. Being alive, time spent while being on dialysis, fatigue and friendliness of staff were the most prominent themes that patients linked to the construct of *dialysis adequacy*.

*Conclusion Adequacy of dialysis* as reported in the literature refers to biochemical outcome measures, most of which are not related with patient relevant outcomes. For patients, adequate dialysis is a dialysis that enables them to spend as much quality time in their life as possible.

## Introduction

Over 2 million people worldwide receive dialysis, and this number is only a fraction of the people that theoretically would need it (1). Although considered a life prolonging technique, the 5-year survival rate of people on dialysis is only 63.3%, and even lower when adjusted for attrition by kidney transplantation (2). Although a small positive evolution in survival can be seen over the years, big improvements are lacking despite numerous studies. Next to the unsatisfactory survival rates, overall quality of life is also poor and the burden of disease high, with lower health related quality of life (HRQOL) (3) indices in hemodialysis patients as compared to the general population (4). The amount of high quality evidence in nephrology is rather low, and harmonization of outcome measures to allow meta-analysis of data across studies to enhance evidence generation, is lacking(5, 6). If we intend to improve the overall care of dialysis patients, it is essential to identify outcomes relevant to all stakeholders so these can be focused on in future research, and to ensure that these outcomes are measured and reported uniformly to allow data aggregation and meta-analysis. The Standardised Outcomes in Nephrology Hemodialysis (SONG-HD) initiative was the first international collaboration that tried to generate this set of ‘core outcomes’ for hemodialysis patients(7, 8). Core outcomes are based on the shared priorities of patients, caregivers and health professionals and are important for decision making. To reduce research waste and increase available evidence, it is essential that all trials include core outcomes as primary or secondary outcome. To generate such a list, the SONG initiative used a validated 5-phase protocol, using several methods and combining qualitative and quantitative data(5). The construct *adequacy of dialysis* was mentioned as an important outcome by patients and health care providers at this stage of the process. However, the meaning of the construct of *adequacy of dialysis* might not be straightforward. The term was coined in the 70s of last century, and gained momentum with the advent of the US National Cooperative Dialysis Study demonstrating greater patient withdrawal from the study and more hospitalizations in the group with the highest blood urea concentrations(9). In this study, treatment failures were associated with fractional small solute clearance, expressed as urea Kt/V(9). Since then, different methods to calculate Kt/V have been introduced(10). More essential, there was increasing acknowledgment that urea per se is not very toxic, and that its behavior during dialysis might not reflect that of potentially more toxic solutes, such as middle molecules or protein bound solutes(11–13). In addition, different authors argued that dialysis adequacy should also cover control of uremic symptoms, extracellular volume and blood pressure, and improve survival and quality of life(14). It can thus be expected that the construct *adequacy of dialysis* as reported in randomized controlled trials or used in the literature, covers a broad scope of substantially different items.

Over the last decade, increasing emphasis has been placed on patient involvement and patient centredness of care. Shared decision making can be an helpful tool to enhance patient centredness, as the technique maximizes the probability that interventions will result in outcomes meaningful to patients(15). To achieve this, it is essential that sufficient quality data on the link between an intervention and the relevant outcomes are available, and that both professionals and patients use a uniform, unambiguous and unequivocal language.

Therefore, the aim of this review is to evaluate the scope and consistency of the term *dialysis adequacy* used in randomised clinical trials and explore in how far this construct as reported in the literature corresponds to what patients envision from it. Therefore, we performed a systematic review of the definitions of the term adequacy of dialysis and dialysis adequacy used in randomised controlled trials. We also performed semi-structured interviews to explore patient’s views and understanding on this construct.

## Patients and Methods

### Systematic review

We searched the Cochrane Central Register of Controlled Trials (CENTRAL) and Pubmed to identify all randomised controlled trails (RCT) up to July 2017 including adult (>18 years) patients on hemodialysis (P) and reporting *dialysis adequacy*, *adequacy* or *adequacy of dialysis* either as outcome or as covariate in the analysis (O). We restricted papers to those in English, Dutch, French or Spanish (Figure 1, PRISM flow chart).

**Figure 1:**
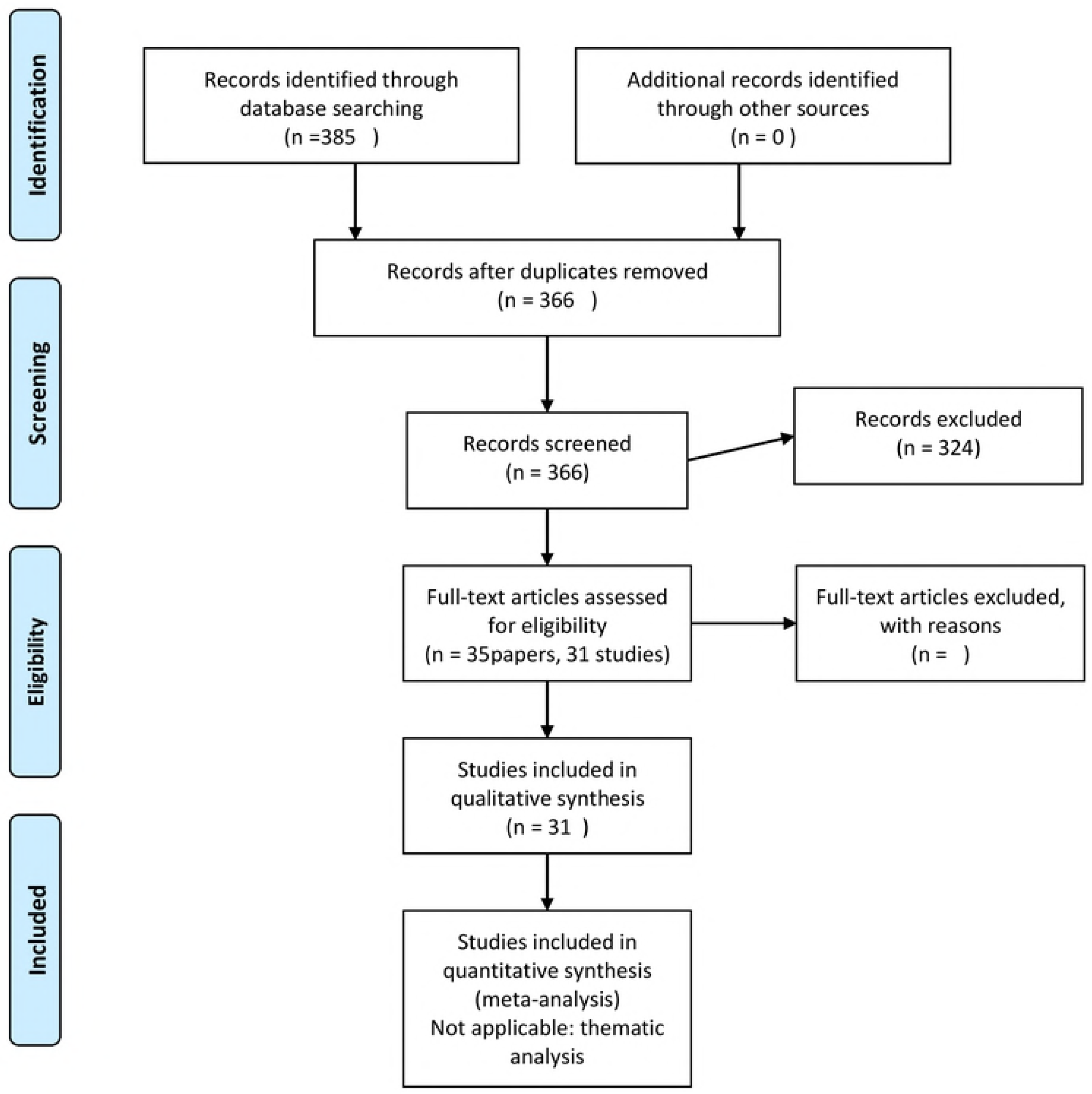
PRISM flow chart of systematic review

**Figure 2:**
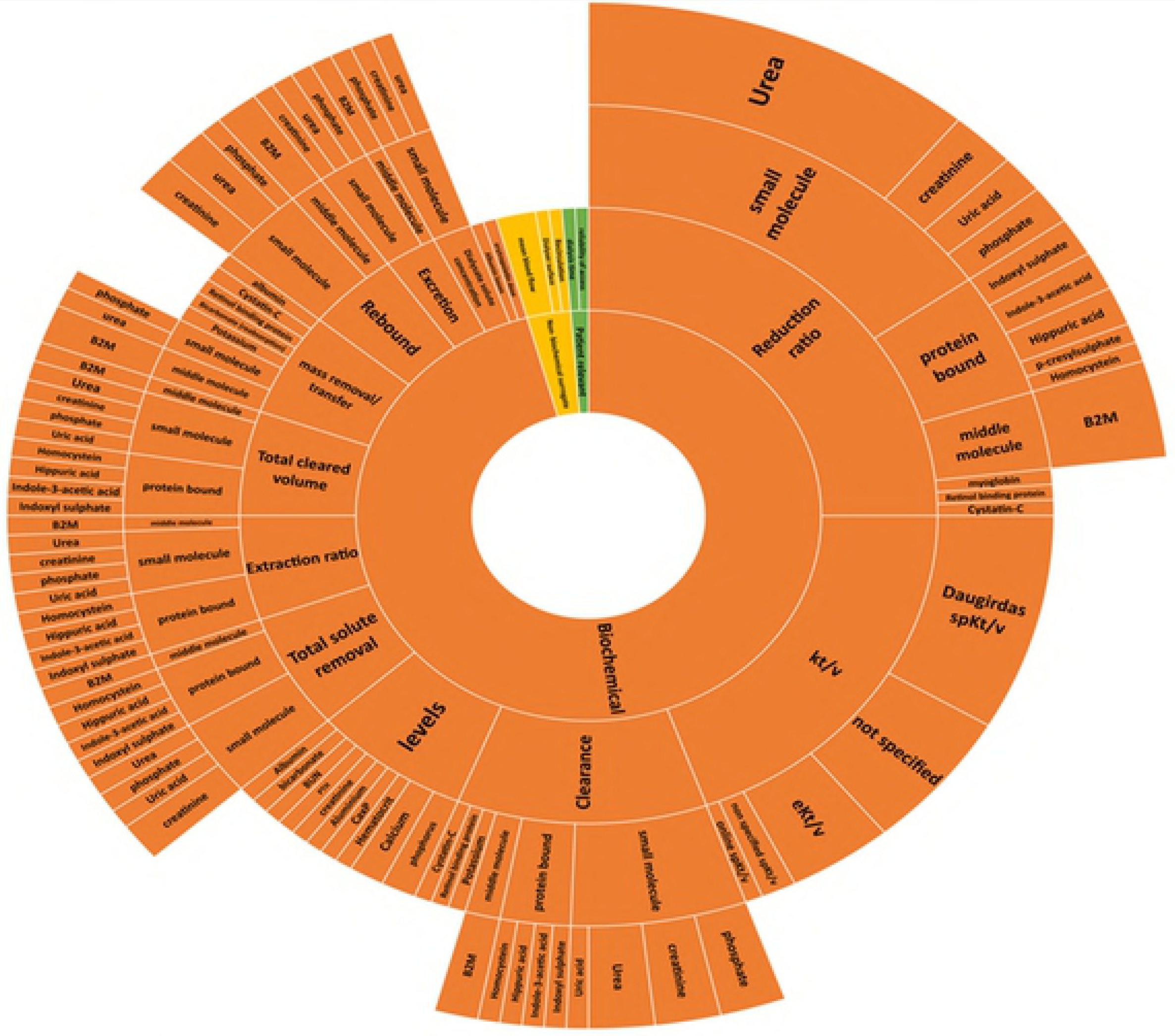
Outcome domains and metrics used to represent adequacy of dialysis in randomized controlled trials

In a first round, we excluded papers not fitting the in‐ and exclusion criteria based on title and abstract. From the remaining studies, we retrieved full texts. We excluded trials that did not include adult patients (18 years of age), studied acute hemodialysis or did not mention *adequacy*, *dialysis adequacy* or *adequacy of dialysis* in the paper.

For each trial, we extracted the following trial characteristics: first author, year of publication, participating countries, sample size, mean age of participants, average study duration, and intervention type. Further, all constructs designated as *dialysis adequacy*, *adequacy of dialysis* or *adequacy* where extracted and counted. As we intended to understand the meaning attributed to *dialysis adequacy*, *adequacy* or *adequacy of dialysis*, we counted all constructs depicted as such in the papers reporting the study, even if this was not the primary or secondary outcome of the study. For example, when a study reported on the association between adequacy of dialysis as defined by Kt/Vurea and quality of life, we counted Kt/Vurea and not quality of life as representing adequacy. For each construct, we extracted domain (e.g. clearance or reduction rate), specific measurement (e.g. urea or creatinine), and timing in relation to the commencement of the trial if applicable. We used studies as measure of unity, so that outcomes of one single study that are reported in different papers on that study are only counted once.

Paper selection and data extraction was done in duplicate (SS and EH). In case of doubt, the item was discussed in the group, and WVB and EN made a final decision.

Per individual item, the total number of times it was used was tabulated and graphically depicted using an Excel work sheet. Items were divided in three major categories: Biochemical (validated or non-validated), non-biochemical surrogate (validated or non-validated) and patient relevant outcomes (relating to clinical items, patient reported outcomes and quality of life). Within each category, when applicable, further distinction was made for different domains (eg clearance, reduction ration, or concentration) and different metrics (eg urea or creatinine) within this domain.

### Semi structured interviews of hemodialysis patients

To check the match of definitions of the construct dialysis adequacy in trials with the understanding of patients, the latter was explored using qualitative methods. Semi-structured interviews were conducted with patients on hemodialysis to ascertain the range of their understanding of the construct *dialysis adequacy*, *adequacy* and *adequacy of dialysis*. The Consolidated Criteria for Reporting Qualitative Health Research (COREQ) were used as reference (16). The study was approved by the Ethical Committee of the Ghent University Hospital (2/2017) and registered with the Belgian registration number B670201731001.

Patients with in center hemodialysis were recruited to participate in an interview. Participants were recruited of different age, gender, social background and education using purposive sampling. Informed consent was obtained from all participants. Inclusion criteria were in center hemodialysis patients age >18years with enough knowledge of French, Dutch or English to create a good contact between interviewer and patient. Patients were excluded when they had important cognitive dysfunction, or were acutely ill.

An interview guide was devised with the following question to prompt the patients to discuss the relevant themes:

1/ Do you know the term “adequacy of dialysis” and what is your understanding of this construct; 2/ what does this construct of “adequacy of dialysis” mean to you personally as a patient; 3/ In literature, “adequacy of diaysis” mainly represents a mathematical (calculated) value based on the amount of urea or one of the other toxins that poison your body if you have severe kidney disease; how do you think about this approach?; 4/ Do you think that “adequacy of dialysis” as represented in literature reflects “good dialysis”? What would “good dialysis mean to you?

Interviews were conducted face-to-face, at the location preferred by the patient. The interview took between 10 minutes and 30 minutes, was audio-taped and transcribed verbatim. Thematic analysis was used to identify the themes reflecting perspectives, beliefs, priorities, and values of patients on hemodialysis regarding *dialysis adequacy* using Nvivo12. Two investigators (SS, MVDV) coded and analyzed the data to develop a first descriptive and analytical identification of themes, which were discussed with WVB and SM (investigator triangulation). Draft results were presented to and discussed with interviewed patients (member checking).

## Results

### Systematic research

We identified 35 articles from 31 trials (table 1, figure 1). With publication dates varying between 1996 and 2016, trials were primarily performed in Europe (13) and North-America (12). Apart from the HEMO-study (5 included trials), there were only two trials in which the number of study participants exceeded 100, and the median number of participants was 39.5 with an interquartile range of 64 (first quartile 15, third quartile 79). Studies were reported as single center trials in 54.8%, and as multicenter investigations in 38.7%, whereas in 3 trials, this was not specified. Only 22% of the studies had a duration of six months or more. All studies combined, the mean age of participants ranged between 55.9 and 74 year.

**Table 1:**
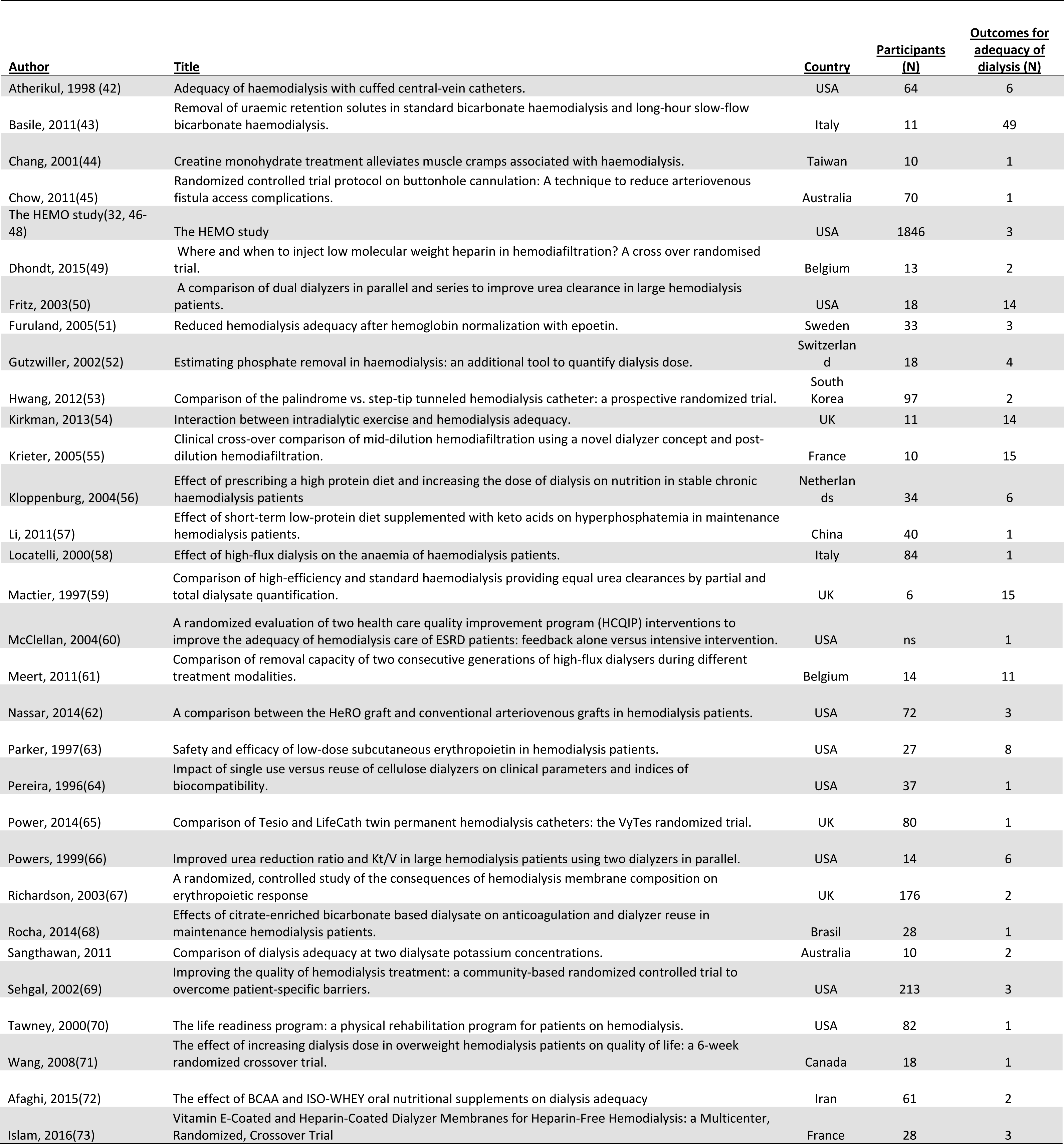
Trial characteristics

The total number of unique markers reported as *adequacy of dialysis* was 98, of which 49 were already provided in one single study. The median number of parameters reported as *adequacy of dialysis* per study was only two. Out of the total of 98 markers, only 5 were not biochemical: mean blood flow, dialyzer surface, recirculation, dialysis time, reliability of access. Except for the blood flow, which was used in three different trials, each of the other patient-relevant and surrogate non-biochemical parameters were only used once.

Every trial used at least one biochemical parameter to represent *adequacy of dialysis*. The most commonly used parameter was Kt/V (87% of the studies). There were several formulas to calculate this however, spKt/V being the most popular method. The second most commonly used parameters was the urea reduction ratio (URR), used by 58% of the trials.

As visible from figure 1, the percentage of the number of trials that reported one of the other specific outcomes declined quickly after these two major indicators. Only clearance of some small molecules (9.6%), the reduction rate of creatinine (9.6%) and beta-2 microglobulin (16.1%) were used in at least three articles. The implementation of other markers was sometimes suggested in the article, but not specifically used within the trial.

### Semi-structured interviews

Patient demographics are represented in table 2.

**Table 2:** Demographics of patients included in the semi-structured interviews

|  |  |
| --- | --- |
| Gender (M/F) | 6/5 |
| Age (years) | 65±12 |
| Time on dialysis (months) | 36±33 |
| Underlying kidney disease | Diabetes (n= 4)<br>Cardiovascular (n=4)<br>Genetic (n=2)<br>Nephrectomy (n=1) |
| Average ultrafiltration (ml/hour) | 410±218 |
| On transplant waiting list (Y/N) | 3/8 |
| Vascular access | AV fistula (n=6)<br>Tunneled catheter (n=5) |

Only 1 out of the 11 patients and explained *adequacy of dialysis* as the degree of removal of waste products. When we reframed the question as ‘good dialysis’, 5 out of 11 patients indicated that dialysis was good simply because they were still alive, and they accepted this treatment in order to live longer.

In almost every interview, patients indicated they did not have a choice but come to dialysis. They could either conform to the suggested treatment, or they would die. Some clearly stated they were happy dialysis existed because it allowed them to live longer. The time spent on dialysis was the price they had to pay. Ways of dealing with the lost time were rationalization and comparison. One patient mentioned that for her, adequacy was the usefulness of dialysis. It allowed her to live a free and normal live in between sessions.

> *“When you poison your own body, I am content this exists so they can help me. I need to come here three times a week but it is something you have to do…if you want to live that is” patient 2, male, 64y*
>
> *“It has to happen, otherwise I die. You don’t have a choice”. Patient 11, male*
>
> *“If there is no other option. You are happy to accept it then”. Patient 9, male, 69y*

Other themes that emerged during the interviews were clustered around 1/organization of their life and time in relation to dialysis; 2/ physical experience of dialysis; 3/ social and external factors; 4/ coping; 5/ dialysis related themes (table 3).

**Table 3:**
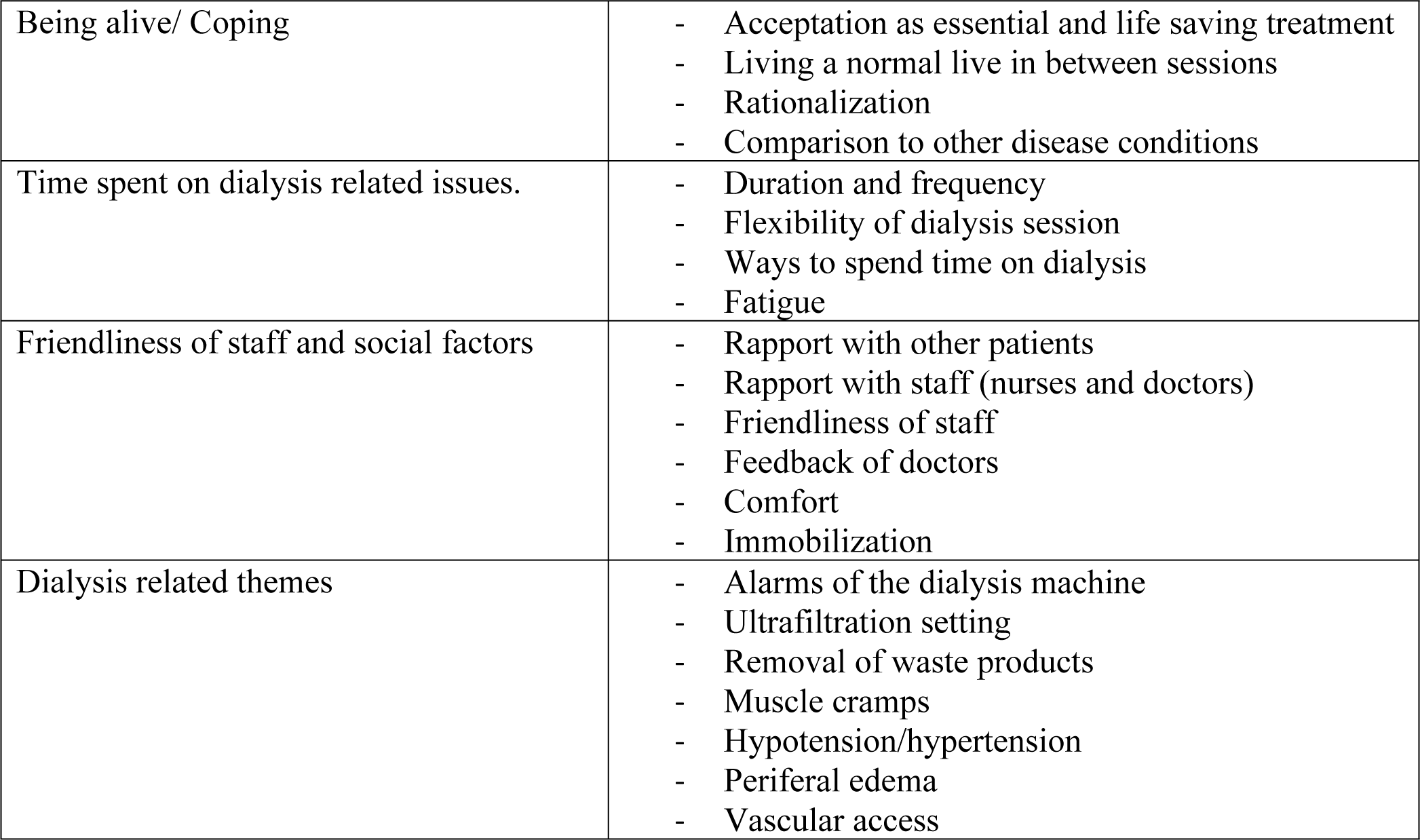
Themes from the semi-structured interviews

The time people spent for their dialysis frequently emerged as a theme. Patients considered longer dialysis sessions as inversely correlated with good dialysis. Although the time spent actually on dialysis was most frequently quoted, also time spent with transport or waiting to be connected were often mentioned. Remarkably, all interviewed patients would accept longer or more dialysis sessions if there was certain proof it would improve their survival or quality of life.

Some patients mentioned flexibility in the timing of the dialysis session as a major aspect of “adequate dialysis”.

> *“There are nurses that complain when I am five or ten minutes late. But don’t forget, I can’t make it on time when there is work.” patient 5, male, 56y*

Importantly, most patients consider also the wash out period and the period with lack of energy after a dialysis session as “wasted time”

> *“One time, I had to come three days in a row. I don’t know the reason anymore but it was too much. I can’t handle it. I was so tired, no energy for anything.” Patient 4, female*

Some patients considered the time on dialysis as an opportunity to rest or sleep so they have more energy in between dialysis sessions.

> *“I come here to rest for four hours. Here I can sleep if I want”, patient 9, male, 69y*

Fatigue was an ambivalent theme in dialysis adequacy. While some people explained they knew dialysis was adequate because they felt very tired before the start of dialysis and less so after, more mentioned fatigue as an unwanted consequence of dialysis, impairing their social life and activities of daily living. Longer dialysis times were often associated with more fatigue, which made that this dialysis was perceived as less good.

> *“When I return from dialysis, I am always tired and have to rest.” Patient 3, female*

Interaction with others during dialysis was seen as an important aspect of good dialysis. Friendliness of staff, mainly nurses, was an often cited theme (5/11), followed by being able to have a chat with fellow patients (4/11). This theme was related to spending time on dialysis, as being able to interact with others made time go by quicker and more meaningful.

> *“The atmosphere and the people. They are all nice people and nurses. For me, that is the important thing. If you cannot chat with anyone or have a laugh…dialysis is not pleasant because it takes so long. If nobody talks, it’s so long.” Patient 6, female, 68y*

Discussions and conflicts with nurses on the other hand could diminish the perception of good dialysis.

> *“Nurses should not be allowed to mess around with you”. Patient 5, male, 56y*

The importance of communication with doctors specifically was not spontaneously mentioned when talking about good dialysis. A good dialysis implied doctors didn’t complain to the patients about laboratory values. However, feedback was appreciated. Patients used it to see if the current diet restrictions were sufficient. When levels of potassium and phosphorus were abnormal, patients stated they had to adjust some eating habits but they did not link it directly with the adequacy of dialysis. For example, although one person would wish for a better cleansing of the blood so she could have more liberty when eating, when there was hyperphosphatemia or hyperkalemia, she would explain it by dietary factors.

> *“A good dialysis is when it everything goes well. When they (doctors) complain, I don’t like it.” Patient 4, female*

The physical changes associated with dialysis could be divided between positive and negative aspects. The development of peripheral edema before and the disappearance of it after dialysis was a way to see that the dialysis works, as were the difference in weight and blood pressure control.

> *“My legs are less swollen” (explaining why hemodialysis is better than peritoneal dialysis) patient 1, female, 72y*

The removal of toxins was only brought up by one person when discussing adequacy of dialysis.

> *“I think a good dialysis mainly means that the clearance of the blood happens in the right way and good way.” Patient 10, female, 38y*

Four people mentioned aspects of the dialysis machine as related with adequacy of dialysis. 2 people followed the amount of ultrafiltration that was set to see if they were strict enough in their fluid intake. A low ultrafiltration need was seen as a good dialysis. Muscle cramps and hypotension were blamed on ultrafiltration, and a dialysis session was considered as good when they could be avoided.

> *“Sometimes my blood pressure drops, then they stop taking fluids away”. Patient 6, female, 68y*

Alarms were seen as a sign of a less good dialysis by 4/11 patients.

> *“A good dialysis is when there are not many alarms. That is a sign that everything goes smoothly.” Patient 7, male, 69y*

Vascular access related topics also emerged during most of the interviews. Although they were never directly linked to adequacy of dialysis, it appeared vascular access has an impact on quality of life of dialysis patients in different ways: fear for problems with access as a lifeline, stress that problems would complicate sessions, capacity for bathing and showers, body disfiguration, prolonged bleeding post-dialysis

> *When you have a catheter, you cannot take a proper bath, or even shower yourself. Patient 6, female, 68 years The bleeding did not stop, and I had to return to the emergency department. Patient 4, female*

## Discussion

The results of the quantitative part of our study indicate that the construct *adequacy of dialysis* is defined substantially differently in different studies. An overwhelming majority of the constructs are based on biochemical surrogate markers, most frequently Kt/V_urea_, which in itself is not uniformly defined however, and urea reduction rate. Patient relevant and surrogate non biochemical parameters were infrequently reported as adequacy of dialysis. The qualitative results of our research demonstrate that the definitions used in randomized controlled trials do not reflect what patients value as *adequate dialysis*. The main themes patients brought forward to represent *adequacy of dialysis* were being alive, time spent on dialysis, fatigue and friendliness of staff and ability for socializing. These themes were in line with previous work(17, 18)

In randomized controlled trials, the removal of small molecules, more specifically Kt/V and the urea reduction rate (URR) are the most commonly used parameters to reflect dialysis adequacy, with 100% of the studies using at least one of them. URR is probably one of the oldest markers used to measure the dose of dialysis(19), but is considered as a rather rough marker of solute removal, as it does not allow to take into account the impact of compartmental behavior of solutes. In a mechanistic post hoc analysis of the randomized National Cooperative Dialysis Study, Kt/V was developed as a measure of prescribed dialysis dose and as a marker to evaluate if an adequate dialysis was delivered. This study coined the terms *adequate dialysis* and *adequacy of dialysis* treatment, basing the validity of this on the observed association of Kt/V with the morbidity of the patient(9).

Since the initial publication, several ways to calculate Kt/V have been developed(20–25). In some jurisdictions, Kt/V is being used as a quality indicator, often even in a summative way. It can be postulated however that adapting the use of Kt/V as an indicator of adequacy of dialysis has several limitations. First and foremost, all associations between Kt/V_urea_ and relevant outcomes are observational, and a proven effect on mortality of changing Kt/V, in whichever form this is calculated, has never been convincingly demonstrated in a randomized controlled trial. Second, focusing on this parameter as the measure for dialysis adequacy ignores the fact that other retention products in uremia are potentially more toxic than urea. In 2012, the European Uremic Toxin Work Group described 88 uremic retention solutes (26, 27), many of them with proven toxicity. Indoxyl sulfate for example is associated with vascular inflammation, endothelial dysfunction and vascular calcification and p-cresylsulphate is a predictor of mortality in patients with varying degrees of kidney impairment (28, 29). As the kinetic behavior of urea and these toxins during dialysis is substantially different from that of urea, Kt/V_urea_ poorly reflects the removal of these uremic toxins during dialysis (13, 30). Reduction ratios of other molecules than urea have also been used in RCTs, but for none of these solutes, indices that take into account compartmental behavior or allow to quantify solute removal, have been developed. Within scientific research, *adequacy of dialysis* indicate substantially different concepts, and in fact poorly reflects removal of the truly toxic substances. The numerous ways of calculating Kt/V_urea_, all reported as *adequacy of dialysis*, make it hard or even impossible to compare results or perform meta-analyses on the topic, further reducing their evidence generating capacity. None of these aforementioned biochemical markers did come up in the patient interviews. Only one patient identified adequacy of dialysis as sufficient removal of waste products. This patient was the youngest and had the highest education level.

For all these reasons, placing Kt/V_urea_ and *adequacy of dialysis* on the same line is cumbersome as the evidence to underpin that better Kt/V_urea_ results in better patient relevant outcomes is largely lacking. For the graphical and hierarchical classification of the outcomes reported as adequacy of dialysis, we used a framework adapted from that used in the SONG-HD initiative(18) and the OMERACT initiative(31). Within the framework, outcome domains are categorized as having vital impact (death), having life impact, and pathophysiological manifestations (surrogate markers). For outcomes labelled as *adequacy of dialysis*, the overwhelming majority of outcomes can be considered as outer tier, as they are purely based on biochemical markers, and have not been validated as being linked in a causal way to patient relevant outcomes. Only a very limited amount of studies used patient relevant outcomes such as quality of life. Neither Kt/V_urea_ nor URR have been validated in a RCT as surrogate markers for higher tier outcomes such as survival or quality of life. In fact, the largest RCT in this regard, the HEMO trial, demonstrates no difference in survival with increasing Kt/V_urea_ (32) Again, the evidence base compiled around the construct of *adequacy of dialysis* is thus very weak, and offers little guidance in decision making because as it lacks studies using robust, meaningful and patient relevant outcomes.

According to the requirements for a quality indicator to be valid for summative monitoring(33), there should be no undesired side effects. This criterium is probably violated when Kt/V_urea_ is used as adequacy parameter. As time and clearance are aspects of this formula, optimizing Kt/V_urea_ values can mean either increasing duration of the dialysis session or increasing the clearance. In the SONG-HD project, dialysis-free time was rated more important by the different stakeholders than relieve of certain uremic symptoms (18, 34). A higher clearance can be achieved by applying higher blood flows or by extending the dialysis duration. The request for higher blood flows can result in a higher number of interventions for vascular access improvement and more hospitalizations. Both of these can have a strong negative impact on quality of life of patients. This might explain why the degree of relation between dose, timing and frequency of dialysis on one hand and quality of life on the other hand is inconsistent in clinical trials (27). Of note, in SONG concerns on vascular access ranked as critically relevant amongst patients(35). In our qualitative analysis, many patients mentioned topics related to their vascular access which can be linked to quality time, such as prolonged bleeding, missed punctures and pain for fistulas and disturbed body image for catheters.

In contrast however to the biochemical definitions of adequacy of dialysis, longer duration of the dialysis session has been associated to improved survival (36, 37). This is an important element to take into the equation during the shared decision process. Patients may however value quality of life as more important than quantity(38). It emerged from our interviews that a good dialysis was considered a dialysis that allowed them as much enjoyable time as possible. The practical and concrete realization of this construct however differed between patients, making it the deepest level of meaning also underlying many other themes. All patients understood that dialysis was a necessity to allow them to life longer, and that they had to forfeit some time for this. This could be either time actually spent on dialysis, or time spent by waiting to be connected or for transport, but also time lost to do preferred activities because of fatigue. The circumstances patients spent their time in during the dialysis session also adds to the lived experience. For some, this is the cozy atmosphere with socializing, laughing and talking with other patients and staff. Other patients rationalize their time, e.g. by reading the newspaper on dialysis rather than at home (“to save time”), or do some administrative work during dialysis. Still others explain they like to take a nap to regain strength for when dialysis is finished. Still other patients loathed the presence of alarms, as they disturbed their impression of a relaxed, homely atmosphere. A good dialysis is according to patients present when all these factors are facilitated by the procedure and staff.

Fatigue often emerged from the interviews as an important symptom associated with adequacy of dialysis. It is often neglected that many patients also loose quality time because of dialysis hangover. Recovery time after dialysis not only reduces quality of life, it is also associated with mortality. For patients, a good dialysis is thus the dialysis that results in as short dialysis recovery time as possible. However, a lower ultrafiltration rate and blood pump speed are associated with shorter recovery time, but also result in more time spent on dialysis, which is by many patients in itself associated with non-good dialysis. It is thus of importance to explain to patients why these interventions are being done. In addition, whereas for some patients, fatigue was induced by dialysis, and good dialysis was thus a dialysis that did not induce fatigue, for others, dialysis washed out their fatigue due to accumulation of toxins. The latter patients saw a positive correlation with the duration or frequency of dialysis and the degree of fatigue, and assessed the adequacy of dialysis on that.

### Strenghts and limitation of the study

We restricted our systematic search to the terms *adequacy of dialysis* and *dialysis adequacy,* to enhance consistency. However, this implied we eliminated articles which only used terms such as ‘dialysis efficacy’ and ‘adequate dialysis dose’. Some might argue that these terms cover the same definition. The reasons we opted to restrict the terms is that we intended to be as objective as possible in our review of the scope of the specific term of *adequacy of dialysis*, limiting the used search terms. As we started our research from a deduction of the SONG conclusion with the particular interest to discover the broad meaning of a construct so highly ranked by patients, using other terms could have provoked other responses in patients. Last, when broadening the scope of used words, we would have created a subjectivity as we would predetermine which words to be valid synonyms for *adequacy of dialysis* and which not.

A potential limitation of the qualitative aspect of this study is related to generalizability due to health literacy and language barriers. As ‘dialysis adequacy’ is an English term, even a B2 word in the CEFR classification (39), we needed to translate it to Dutch for patients to understand it. We did this by applying forward and backward translations using different translators for both directions. Whereas this translation problem can be seen as a limitation of our current research, it would be interested to repeat the semi-structured interviews with both native as non-native speaking patients, to clarify whether the problem is really a language problem (so meaning is changed during translation) or rather a conceptual problem (patients do really consider different items under the construct than researchers do). The interviews strongly suggest that patients intrinsically do attribute different meanings to the construct *adequacy of dialysis,* and that this difference in interpretation is not, at least not solely, induced by imperfect translation.

Although not formally measured, we can presume that health literacy plays an important role in the appreciation of adequacy of dialysis (40). Knowing the working mechanism of a dialysis and understanding the long term consequences of high amounts of toxins and fluid overload may make people focus more on items related with the technical aspects of the dialysis session itself. We could therefore assume the results would be different in for example a home dialysis population. Also the age of the patient might be of importance. In older or frail patients, healthcare workers probably focus more on quality of life as there are less QALYs to gain with stricter regulations of for example phosphate levels (41). This will make communication different than in younger patients, being less rigid and less focused on long term outcomes.

As we were primarily interested in the potential array of themes that emerged when patients are confronted with the construct *adequacy of dialysis* we chose for thematic analysis and purposefully interviewed a diversified patient population in terms of age, gender, dialysis vintage and hemodialysis modality. It appeared that every individual patient has his/her own attributes to this term, so we did not reach saturation. However, most authors consider data saturation not as indispensable for thematic analysis. In order to gain more insights in the deeper meaning patients convey to the construct *adequacy of dialysis*, a grounded theory approach will be set up in the future. To expand generalizability of our findings, a mixed method approach using questionnaires based on our themes could be used to quantify their prevalence in a broad dialysis population.

### Conclusion

The meaning of the construct *dialysis adequacy* as reported in randomized controlled trials varies between and within researcher groups and patients. None of the definitions patients attribute to adequacy of dialysis is purely biochemical as is nearly uniformly the case when the construct is used in the literature. For patients, the construct rather includes “good enough dialysis to allow a sufficient time with quality of life”. This is an important notion as most of the strategies to improve biochemical definitions of adequacy of dialysis are thus in conflict with what patients appreciate and value. In order to improve shared decision making, evidence needs to be generated to link both worlds together.

